# Context versus aiming in motor learning when both feedforward and feedback control processes are engaged

**DOI:** 10.1101/2023.11.28.569130

**Authors:** Matthew J. Crossley, Christopher L. Hewitson, David M. Kaplan

## Abstract

Theories of human motor learning commonly assume that movement plans are adjusted in response to the precision of sensory feedback received regarding their success. However, support for this assumption has mainly come from experiments that limit feedback correction during an ongoing movement. In contrast, we have recently shown that when this restriction is relaxed, and both within-movement and between-movement corrections can occur, movement plans undergo large and abrupt changes that are strongly correlated with the degree of sensory uncertainty present on the previous trial and are insensitive to the magnitude and direction of recently experienced movement errors. A class of models in which sensory uncertainty influences an aiming process with no retention from one trial to the next best accounted for these data. Here, we examine an alternative possibility that sensory uncertainty acts as a contextual cue to shunt motor learning and control to one of many context-specific internal models. Although both aiming and context models provide good fits for our data, the aiming model performed best.

**Author summary:** A large body of literature shows that sensory uncertainty inversely scales the degree of error-driven corrections made to motor plans from one trial to the next. However, by limiting sensory feedback to the endpoint of movements, these studies prevent corrections from taking place during the movement. We have recently shown that when such corrections are permitted, sensory uncertainty punctuates between-trial movement corrections with abrupt changes that closely track the degree of sensory uncertainty but are insensitive to the magnitude and direction of recently experienced movement error. Here, we ask whether this pattern of behaviour is more consistent with sensory uncertainty driving changes in an aiming process or context-specific motor learning.

## Introduction

We have recently shown that when both within-movement and between-movement error corrections are permitted, movement plans undergo large and abrupt changes that are strongly correlated with the degree of sensory uncertainty present on the previous trial and are insensitive to the magnitude and direction of movement errors [1]. These results diverge from existing studies which have all shown that the adaptation of movement plans inversely scales with the level of uncertainty present in the sensory feedback [2–8]. Instead, our results appear most consistent with a class of models that assumes sensory uncertainty influences an *aiming* process that contains no retention from one trial to the next and is therefore completely determined by the sensory uncertainty experienced on the previous trial.

However, our previous analysis overlooked an important class of models. In particular, rather than triggering discrete aiming, sensory uncertainty may instead serve as a powerful contextual cue, shunting learning and motor output into one of potentially many context-specific internal states. We explore this possibility here. Here, we extended our previous analysis by attempting to interpret our results through the lens of contextual inference. According to several influential theories, motor learning is exquisitely context-specific [9–11]. Although varied in their implementation and specific assumptions, these theories all roughly assume that the brain maintains many internal models for different context-appropriate sensorimotor mappings. From this perspective, it is possible that different sensory uncertainty conditions engage different context-specific internal models.

## Results

Fig 1a shows group-averaged initial movement vectors observed in human participants from our previous report [1]. This panel clearly shows that cahnges in initial movement vectors across trials (e.g., as would be driven by feedforward motor adaptation) undergo large and abrupt changes that are strongly correlated with the degree of sensory uncertainty present on the previous trial and are largely insensitive to the magnitude and direction of movement errors.

**Fig 1.**
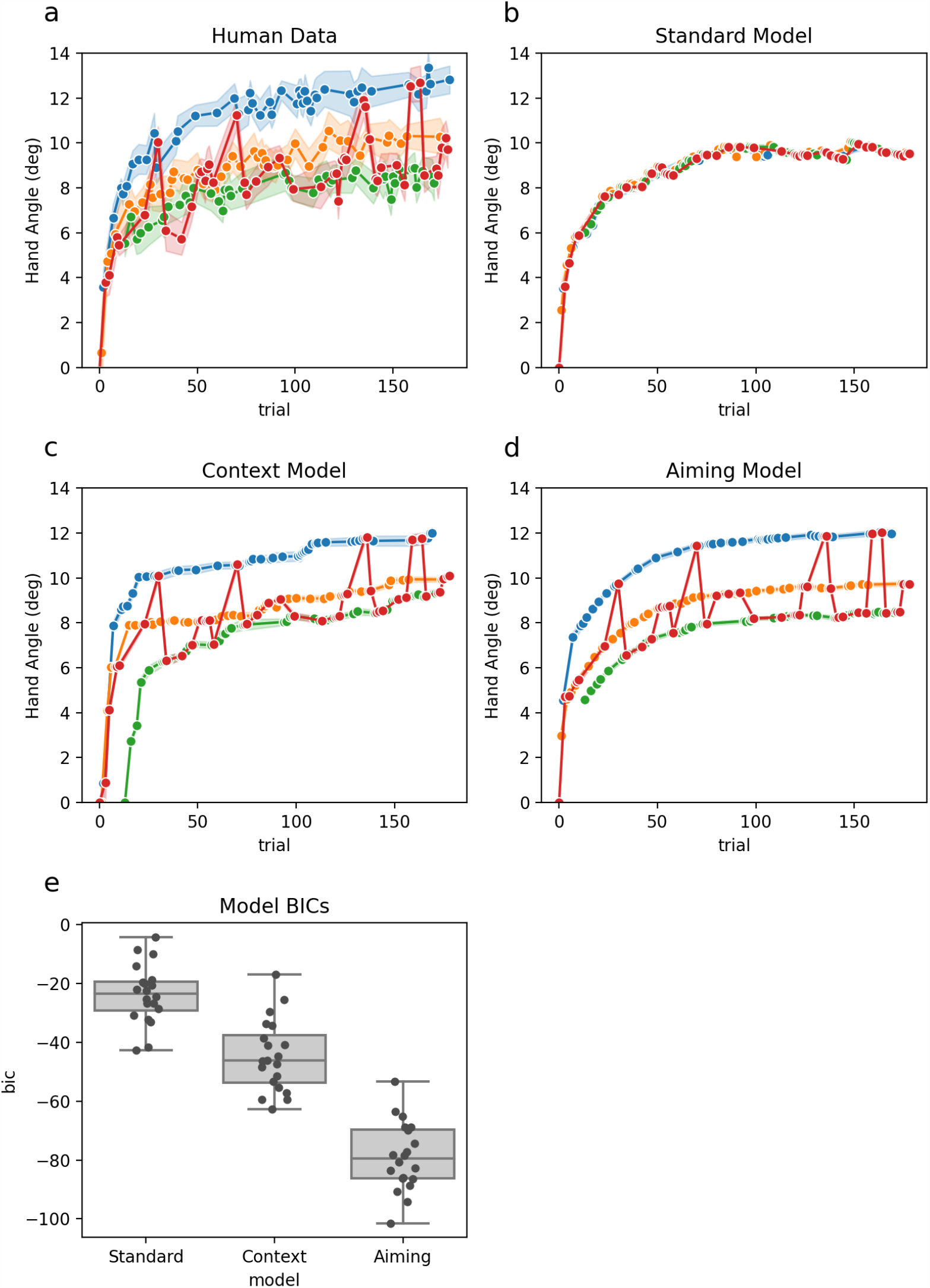
**a:** Mean initial movement vector across participants per trial. **b:** Results predicted by the standard model. **c:** Results predicted by the context model. **d:** Resuts predicted by the aiming model. **e:** BIC scores for all three models. Line and markers in panels a–d are color-coded according to the sensory uncertainty present on trial *t* − 1. Error bands are 95% CI.

The *standard model* shown in Fig 1b fails to predict any striation between different levels of sensory uncertainty. Both the *context model* shown in Fig 1c and the *aiming model* shown in Fig 1d capture the majour striation between sensory uncertainty levels and offer plausible accounts of the behavioural data. However, due to the *context model* mispredicting the first few trials, the *aiming model* was clearly preferred by BIC in nearly every case.

## Discussion

Classic models of motor learning assume that the amount of adaptation that will occur in response to a movement error is proportional to the magnitude of the error [2, 12–17] and inversely proportional to the uncertainty of the sensory signal conveying the error [4, 5, 18–20]. We have shown both here and in our previous work [1] that this *standard model* does not recapitulate behavioural data from our novel paradigm which was chacterized by large, abrupt, and frequent changes in initial movement vectors across trials.

On the other hand, an *aiming model* assuming that different levels of sensory uncertainty trigger different aiming strategies easily succeeded. This model is consistent with the idea that sensory uncertainty triggers an explicit aiming strategy [21–24] though the aiming process need not strictly be explicit. At first glance the *aiming model* seems to offer cogent explanation of an otherwise rather puzzling pattern of behavioural data. On closer inspection, it is unclear why high uncertainty trials would prompt participants to abandon a previously successful explicit strategy, only to suddenly revert to the same strategy just a few trials later. Additionally, it is difficult to imagine an aiming process that would act consistently and with so little variation across participants as is implied by our data.

Here, we extended our previous analysis by attempting to interpret our results through the lens of contextual inference. According to several influential theories, motor learning is exquisitely context-specific [9–11]. Though varied in their implementation and specific assumptions, these theories roughly assume that the brain maintains many internal models that learn context-appropriate sensorimotor mappings. From this perspective, it is possible that different sensory uncertainty conditions engage different context-specific internal models. Here, we showed that such a *context model* can indeed recapitulate the major qualitative aspects of our behavioural data.

The *context model* may be preferred because it elegantly avoids the previoulsy mentioned puzzles attached to the *aiming model*. On the other hand, the *aiming model* significantly outperformed the *context model* by formal model comparison metrics. Furthermore, the processes posited by each model is well supported by a large preexisting literature. Ultimately, we must leave the resolution of this puzzle to future research.

## Conclusion

Standard models assume that the degree of between-movement motor correction is inversely proprorional to the uncertainty of the sensory signal conveying movement errors. These models fail only if (1) sensory uncertainty varies trial-by-trial and (2) the experimental paradigm promotes within-movement corrections. We propose two viable alternatives. The *aiming model* assumes that different levels of sensory uncertainty trigger different aiming strategies, and the *context model* assumes that different levels of sensory uncertainty trigger different context-specific adaptive states. Of these, the *aiming model* provides the best account of our existing data.

## Materials and methods

### Behavioral experiment

Participants performed reaches with their dominant (right) hand from a starting position located at the center of the workspace to a single target located straight ahead. The experiment included a brief baseline phase in which veridical feedback was provided, an adaptation phase in which perturbed feedback was provided, and a washout phase in which feedback was withheld entirely. Here we only consider data from the adaptation phase. During the adaptation phase, feedback was provided only at movement midpoint and endpoint and was displayed at one of four visual uncertainty levels (*σ*_*L*_, *σ*_*M*_, *σ*_*H*_, *σ*_*∞*_). Across trials, both the sequence of perturbations and the sequence of sensory uncertainties was matched across participants. The data from the behavioral experiment has been previously reported in Experiment 2 of our recent publication [1]. Please consult that paper for further methodological detail.

### State-space models

We recently developed several models which fall into two classes of assumptions regarding how sensory uncertainty influences motor learning [1]. The first class assumed that sensory uncertainty influences the updating of the adaptive learning process and includes classic *error-scaling* models. The second class assumed that sensory uncertainty influences an aiming process that contains no retention from one trial to the next and is therefore completely determined by the sensory uncertainty experienced on the previous trial. The *aiming* models performed better overall than any other model we explored. Here, we consider the possibility that, rather than driving an aiming process, sensory uncertainty may instead serve as a powerful context cue which acts to shunt learning and output into one of potentially many context-specific internal states.

The following model descriptions favor simplicity over completeness. In particular, each model includes an internal state variable that controls within-movement error correction, yet below we only specify the aspects of the model pertaining to feedforward motor planning. Moreover, while updates to the internal state governing feedforward motor plans (i.e., between-movement error correction) depends on the sensed error both at movement midpoint and also at movement endpoint, here we refer only to movement error. This simplification is warranted because we consider only data from a condition in which the uncertainty of sensory feedback is matched at these two time points. For full details on these aspects of the models please consult our earlier work [1].

#### The Standard model

The *standard model* assumes that the uncertainty of sensory feedback influences the update of an internal state variable by acting as a gain on the learning rate:

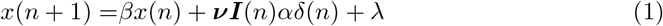

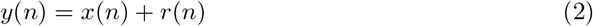

Here, *n* is the current trial, *δ* is the sensed movement error, *y* is the motor output, *r* is the imposed rotation, and *x* is an internal state variable. The *β* parameter is a retention rate that describes how much is retained from the value of the state at the previous trial, *α* is a learning rate that describes how quickly states are updated in response to errors, and *λ* is a constant bias. The ***ν*** = [*ν*_0_, *ν*_*M*_, *ν*_*H*_, *ν*_*∞*_] variable is a row vector of free parameters (one value for each level of sensory uncertainty), ***I*** is a column vector that indicates the uncertainty of sensory feedback that was present on trial *n*. The classic idea that sesnory uncertainty inversely scales the magnitude of the error-dtiven update would be captured if *ν*_0_ *> ν*_*M*_ *> ν*_*H*_ *> ν*_*∞*_.

#### The context model

The *context model* assumes that different levels of sensory uncertainty act as context cues which shunt motor learning and output to different context-specific internal models:

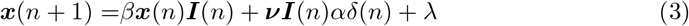

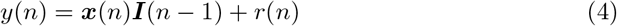

Here, ***x*** = [*x*_0_, *x*_*M*_, *x*_*H*_, *x*_*∞*_] is a row vector of context-specific internal state variables (one value for each level of sensory uncertainty). All other parameters and nomenclature are identical to those described above for the *standard model*. We used single global values for *α, β*, and *γ* (i.e., there were 3 parameters in total, not 3 per context). Note that the the ***I***(*n* − 1) in the equation for *y*(*n*) says that the sensory uncertainty experienced on trial *n* − 1 dictates which context-specific internal state that will drive motor output on trial *n*.

#### The aiming model

The *aiming model* assumes that the uncertainty of sensory feedback directly influences the feedforward motor output:

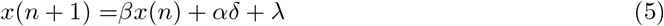

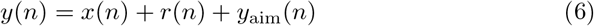

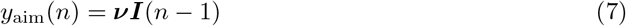

All parameters and nomenclature are identical to those described above for *standard model*. Note that the the ***I***(*n* − 1) in the equation for *y*_aim_(*n*) says that the sensory uncertainty experienced on trial *n* − 1 dictates the aiming on trial *n*.

#### Parameter estimation and model comparison

For each model, we obtained best-fitting parameter estimates on a per subject basis by minimising the sum of squared error difference between the observed and predicted midpoint and endpoint hand angles. We used the differential evolution optimization [25] method implemented in *SciPy* [26] to find the parameter values that minimised this metric. Table 1 shows the bounds for each parameter to which this optimization was constrained. We then computed the Bayesian Information Criterion (*BIC*) as *BIC* = *nlog*(1 − *R*^2^) + *klog*(*n*) where *k* is the number of model parameters, *n* is the number of observations, and *R*^2^ is the proportion of variance explained by the optimised model. Models with lower *BIC* value are preferred [27].

**Table 1.**
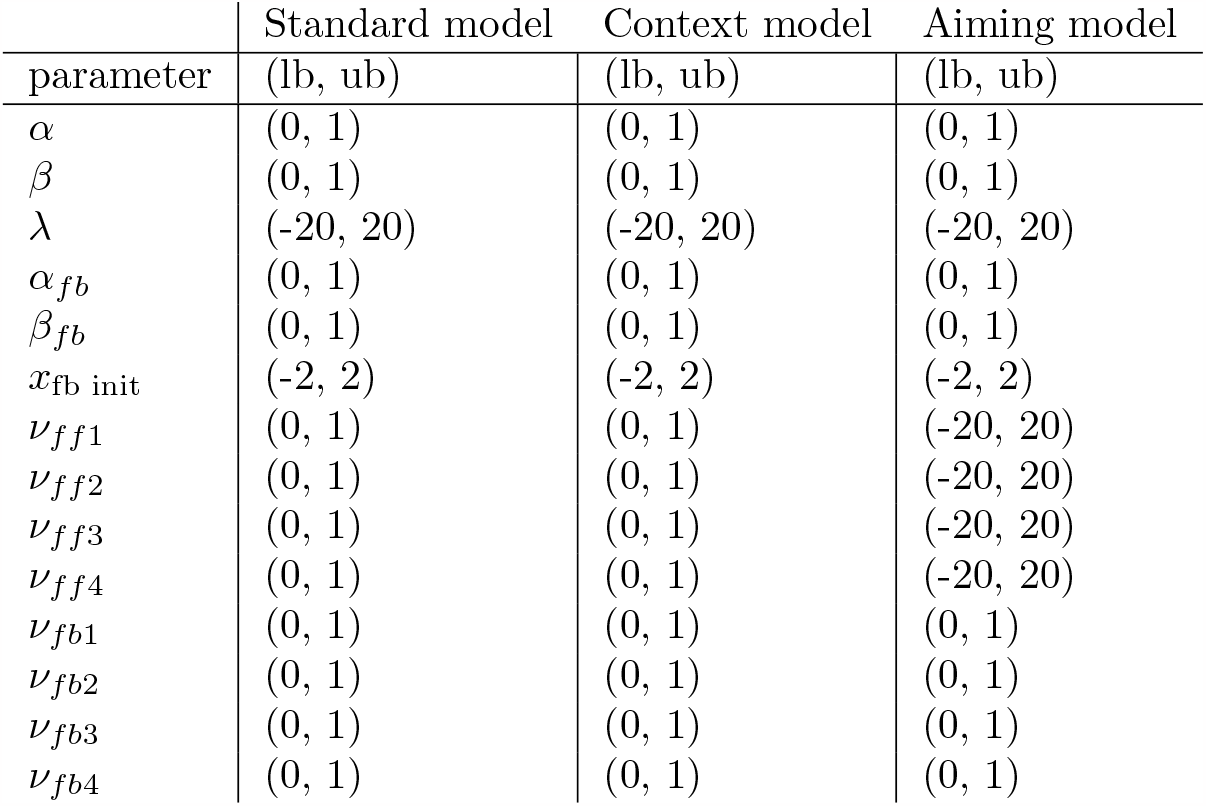
State-space model parameter bounds. Lower and upper values are indicated by (lb, ub), respectively. For details on *α*_*fb*_, *β*_*fb*_, *x*_fb init_, *ν*_*fb*1_, *ν*_*fb*2_, *ν*_*fb*3_, and *ν*_*fb*4_ please see [1]

## Code Accessibility

Data and analysis code can be accessed at: https://github.com/crossley/sensory_uncertainty_fffb_fffb_context_vs_aim

## References

1. Hewitson CL, Kaplan DM, Crossley MJ. Error-independent effect of sensory uncertainty on motor learning when both feedforward and feedback control processes are engaged. PLoS computational biology. 2023;19(9):e1010526.

2. Scheidt RA, Dingwell JB, Mussa-Ivaldi FA. Learning to Move Amid Uncertainty. Journal of Neurophysiology. 2001;86(2):971–985. doi:10.1152/jn.2001.86.2.971.

3. Baddeley RJ, Ingram HA, Miall RC. System Identification Applied to a Visuomotor Task: Near-Optimal Human Performance in a Noisy Changing Task. The Journal of Neuroscience. 2003;23(7):3066–3075. doi:10.1523/JNEUROSCI.23-07-03066.2003.

4. Burge J, Ernst MO, Banks MS. The statistical determinants of adaptation rate in human reaching. Journal of Vision. 2008;8(4):20. doi:10.1167/8.4.20.

5. Wei K. Uncertainty of feedback and state estimation determines the speed of motor adaptation. Frontiers in Computational Neuroscience. 2010;doi:10.3389/fncom.2010.00011.

6. Verstynen T, Sabes PN. How Each Movement Changes the Next: An Experimental and Theoretical Study of Fast Adaptive Priors in Reaching. Journal of Neuroscience. 2011;31(27):10050–10059. doi:10.1523/JNEUROSCI.6525-10.2011.

7. Fernandes HL, Stevenson IH, Kording KP. Generalization of Stochastic Visuomotor Rotations. PLoS ONE. 2012;7(8):e43016. doi:10.1371/journal.pone.0043016.

8. Fernandes HL, Stevenson IH, Vilares I, Kording KP. The generalization of prior uncertainty during reaching. Journal of Neuroscience. 2014;34(34):11470–11484.

9. Heald JB, Lengyel M, Wolpert DM. Contextual inference underlies the learning of sensorimotor repertoires. Nature. 2021;600(7889):489–493. doi:10.1038/s41586-021-04129-3.

10. Lee JY, Schweighofer N. Dual Adaptation Supports a Parallel Architecture of Motor Memory. Journal of Neuroscience. 2009;29(33):10396–10404. doi:10.1523/JNEUROSCI.1294-09.2009.

11. Wolpert DM, Kawato M. Multiple paired forward and inverse models for motor control. Neural Networks. 1998;11(7-8):1317–1329. doi:10.1016/S0893-6080(98)00066-5.

12. Kawato M, Furukawa K, Suzuki R. A hierarchical neural-network model for control and learning of voluntary movement. Biological cybernetics. 1987;57(3):169–185.

13. Wolpert DM, Miall RC, Kawato M. Internal models in the cerebellum. Trends in Cognitive Sciences. 1998;2(9):338–347. doi:10.1016/S1364-6613(98)01221-2.

14. Thoroughman KA, Shadmehr R. Learning of action through adaptive combination of motor primitives. Nature. 2000;407(6805):742–747. doi:10.1038/35037588.

15. Cheng S, Sabes PN. Modeling Sensorimotor Learning with Linear Dynamical Systems. Neural Computation. 2006;18:760–793.

16. Tanaka H, Sejnowski TJ, Krakauer JW. Adaptation to visuomotor rotation through interaction between posterior parietal and motor cortical areas. Journal of neurophysiology. 2009;102(5):2921–2932.

17. Tanaka H, Krakauer JW, Sejnowski TJ. Generalization and multirate models of motor adaptation. Neural computation. 2012;24(4):939–966.

18. Korenberg AT, Ghahramani Z. A Bayesian view of motor adaptation. Current Psychology of Cognition. 2002;21(4/5):537–564.

19. Wei K, Kording K. Relevance of error: what drives motor adaptation? Journal of neurophysiology. 2009;101(2):655–664.

20. Tsay JS, Avraham G, Kim HE, Parvin DE, Wang Z, Ivry RB. The effect of visual uncertainty on implicit motor adaptation. Journal of neurophysiology. 2021;125(1):12–22.

21. Taylor JA, Ivry RB. Flexible Cognitive Strategies during Motor Learning. PLoS Computational Biology. 2011;7(3):e1001096. doi:10.1371/journal.pcbi.1001096.

22. Taylor JA, Krakauer JW, Ivry RB. Explicit and Implicit Contributions to Learning in a Sensorimotor Adaptation Task. Journal of Neuroscience. 2014;34(8):3023–3032. doi:10.1523/JNEUROSCI.3619-13.2014.

23. McDougle SD, Ivry RB, Taylor JA. Taking Aim at the Cognitive Side of Learning in Sensorimotor Adaptation Tasks. Trends in Cognitive Sciences. 2016;20(7):535–544. doi:10.1016/j.tics.2016.05.002.

24. Tsay JS, Haith AM, Ivry RB, Kim HE. Interactions between sensory prediction error and task error during implicit motor learning. PLoS computational biology. 2022;18(3):e1010005.

25. Storn R, Price K. Differential evolution–a simple and efficient heuristic for global optimization over continuous spaces. Journal of global optimization. 1997;11(4):341–359.

26. Virtanen P, Gommers R, Oliphant TE, Haberland M, Reddy T, Cournapeau D, et al. SciPy 1.0: fundamental algorithms for scientific computing in Python. Nature methods. 2020;17(3):261–272.

27. Kass RE, Raftery AE. Bayes factors. Journal of the american statistical association. 1995;90(430):773–795.

